# Elements of Olfactory Intelligence in *Drosophila*

**DOI:** 10.64898/2026.01.05.697602

**Authors:** Aurel A. Lazar, Yiyin Zhou

## Abstract

The ability to make the world of odorants intelligible is a key capability of the *Drosophila* olfactory system that we shall call olfactory intelligence. Characterizing the functional logic of the *Drosophila* early olfactory system that makes the natural world of odorants intelligible is a major challenge in neuroscience.

Starting by modeling the space of odorants using constructs of both *semantic* and *syntactic* information, we establish that the Antenna Lobe and Mushroom Body Calyx first decompose the confounding representation of the Antenna into a concentration independent odorant semantics information and the ON-OFF timing of the syntactic information. Subsequently the two streams of information are integrated and produce a novel time and rank-based representation of Kenyon Cell outputs, called the marked first spike sequence code. In conjunction with a novel distance measure for the spike sequence code, we demonstrate that the rank-based representation supports accurate classification of ON-OFF odorant semantics. Computationally, these elements of olfactory intelligence are realized by a class of *differential* divisive normalization processors modeling the feedback circuits in a causal chain of stages including the Antenna, Antennal Lobe and the Mushroom Body Calyx. Consequently, the early olfactory system makes the natural world of odorants intelligible.

## 1 Introduction

*Drosophila* olfactory circuits sense and process a wide range of information streams arising in many environmental niches [1]. The odorant spaces associated with these niches contain, among others, identifiable olfactory objects of interest. The early olfactory system (Figure 1 (top)) evolved to identify and characterize in detail the olfactory object features of their respective sensory world. But what is the structure and what are the identifiable objects in these spaces? To what extent are these objects similar and/or different? Formally characterizing the (i) objects of interest in the olfactory space of *Drosophila’s* natural sensory world, and the (ii) functional logic of the brain circuits extracting the odorant object features, are major challenges in neuroscience.

**Figure 1.**
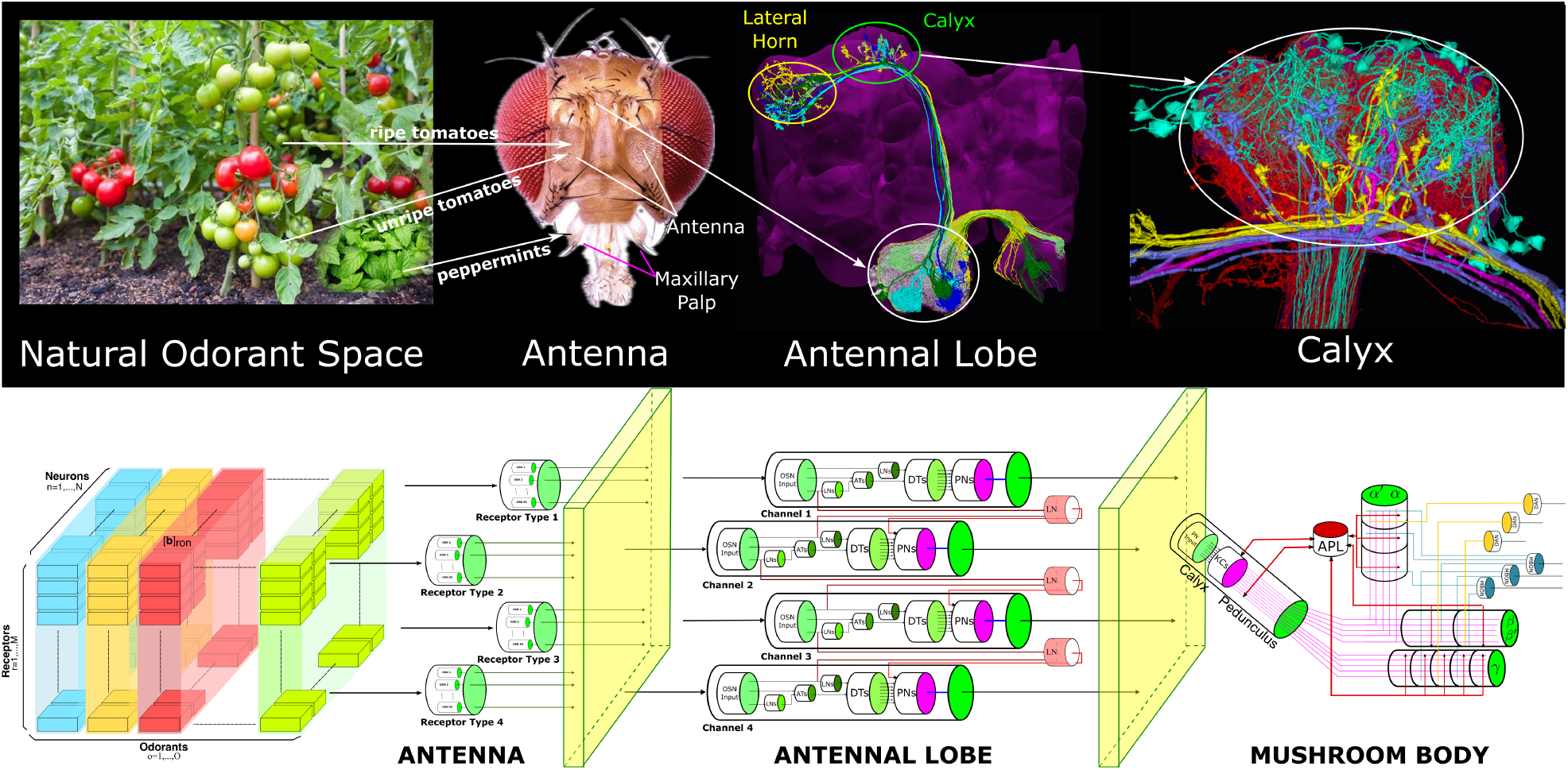
(top) Odorant processing pathways of the Early Olfactory System of the fruit fly. (bottom) Structural pathways of the Early Olfactory System of the fruit fly. Only the right hand side of the EOS is shown.

Central to our interests is the intelligent discrimination of odorant identity as an object of the olfactory world. It requires high accuracy in processing of possibly known and/or novel odorants encountered in novel settings/environments. Thus, in agreement with the broad outline in [2] we argue that taking the olfactory world to consist of intelligible odorant objects is an expression of intelligence and investigate how the *Drosophila* finds the olfactory world intelligible.

### 1.1 Semantic and Syntactic Odorant Information Flows

The three stages of the Early Olfactory System (EOS) of the fruit fly brain that we will be focussing on are the Antenna (AN), the Antennal Lobe (AL) and the Mushroom Body (MB) Calyx.

As shown in Figure 1(top), the natural odorant environment (see Figure 1(top, column 1)) is first sensed by the Olfactory Sensory Neurons (OSNs) whose cilia are hosted in sensilla located on the Antennae and Maxillary Palps (shown in Figure 1(top, column 2)) [3]. Olfactory transduction takes place on the dendrites (cilia) of the OSNs. The chemical odorants are transduced into electrical action potentials [4].

OSNs expressing the same olfactory receptor (OR) type project into an individual glomerulus of the Antennal Lobe (AL) (see Figure 1, top column 3) [5]. The dendrites of uniglomerular Projection Neurons (PNs) innervate a single glomerulus, receiving input from OSNs expressing the same OR-type. The PNs then project into the MB Calyx and provide inputs to Kenyon Cells (KCs) [5]. In the MB Calyx, KC dendrites receive feedback from an Anterior Paired Lateral (APL) neuron with inputs from all the KCs (see Figure 1, top column 4) [6, 7].

It is critical to note that two different types of information flows are present during the chemical to electrical transduction process.

The first information flow is due to the chemical identity information of the odorant molecules acquired through a binding and dissociation process by the OSN expressed olfactory receptors. Each receptor type has a different binding and dissociation rate for each type of odorant molecules [8, 9]. Since this type of information characterizes the identity of an odorant, we call it *semantic information* [10]. Semantic information relates to “meaning” [11]. An example of “semantics” in olfaction is the “smell of a rose”. Acetone might be perceived to be “aversive” or “attractive”.

The second information flow is due to the time-dependent concentration waveform. A higher concentration amplitude typically leads to a stronger OSN response [8, 9]. We refer to the odorant concentration waveform as *syntactic information* [10]. Syntactic information (also known as Shannon information [11]) characterizes the concentration amplitude of the odorants of the biological environment.

While Shannon information is rather well characterized, semantic information has been extremely difficult to formalize. Simply put, there is a dearth of research in modeling, characterizing and processing semantic information in neuroscience. A key observation regarding the very first stage of olfactory sensing is that, as we will show in Section 2.3, olfactory transduction is a complex process where odorant semantic information and syntactic information are multiplicatively coupled in individual OSNs. In other words, the early olfactory system of the fruit fly encodes the odorant object identity (semantic information) and the odorant concentration waveform (syntactic information) into a combinatorial neural code [8, 9]. Therefore, in order to distinguish in time the presence of an odorant, the two odorant attributes must be untangled by the downstream circuit.

This calls for (i) formally modeling odorant stimuli in terms of their semantic and syntactic information content, and for (ii) exploring a new class of processors, called Intelligent Olfactory Processors (IOPs), that gradually shift in the early stages of the olfactory system to processing olfactory semantic and syntactic information in parallel. Concluding, starting from multiplicatively coupled semantic and syntactic information flows, IOPs provide an ON-OFF representation of odorant semantics at the input of the MB olfactory associative memory circuits that support a high degree of semantic specificity (i.e., separation between different odorant identities).

In this article, we describe our approach to address these elements of olfactory intelligence in the early olfactory system of the fruit fly.

### 1.2 Causal Odorant Processing Pathways in the *Drosophila* EOS

The *Drosophila* EOS depicted on the top of Figure 1 is clearly a causal system. Thus, *Drosophila* EOS processing models have to critically abide by causality; time plays a critical role in such models. Odorant object identities may not intrinsically carry a time dimension, and the temporal variability is often neglected or avoided in many accounts of olfactory processing models. Causality also implies that the information flows are processed stage by stage starting from the AN periphery. A characterization in the time doamin of the output of each processing stage in Figure 1 is critical for determining the information flow to the next processing stage.

The schematics of the odorant space model and the causal odorant processing pathways of the EOS are shown in Figure 1(bottom, columns 1, and 2 to 4, respectively). The AN circuit (Figure 1, bottom column 2) consists of parallel OSNs that are randomly distributed across the surface of the Maxillary Palp and AN. The OSNs are depicted in groups based on the olfactory receptors that they express. The OSNs provide input to the AL (shown in Figure 1, bottom column 3). The multiplicatively coupled olfactory signal at the output of the AN is mapped into a code in the spike domain carried by the PNs. The PNs feed into Kenyon Cells (KCs) that are the primary outputs of the MB Calyx (see Figure 1, bottom column 4) and that project to the 15 Mushroom Body compartments where associative learning takes places [12].

The information present in the code underlying the KC spike trains, is a key challenge towards elucidating the functional logic of olfactory associative learning. How is the odor information progressively transformed across different neuropil stages (Figure 1 top row) into formats that lead to a robust classification of ON-OFF odorant semantics?

To address this challenge we closely follow the workflow, first described in [13], for discovering the functional logic of *Drosophila* brain circuits.

### 1.3 Manuscript Organization

In Section 2, a novel Odorant Encoding Machine (OEM) modeling odorant processing in the EOS is formulated stage by stage, leading to an understanding of its functional logic. The first element of olfactory intelligence is discussed in Section 2.1, where the space of pure odorants is characterized in terms of semantic and syntactic information. The second element of olfactory intelligence is discussed in Sections 2.2-2.5, where we detail some of the schematics of the OEM and the Intelligent Olfactory Processors (IOPs) in the AN, the AL and the MB Calyx. The IOPs operate in parallel on olfactory semantic and syntactic information flows. In Section 3, we advance a rank-based distance measure to evaluate the semantic specificity of the marked first spike sequence code under different circuit configurations. An overall discussion of the key elements of olfactory intelligence in the *Drosophila*’s EOS is presented in Section 4.

## 2 The Functional Logic of the *Drosophila* Early Olfactory System

In this section, we describe an executable model of the EOS schematically outlined in Figure 1 (top: morphology, bottom: graph structural pathways).

Section 2 is organized as follows. A model of the space of odorants that explicitly takes into account the odorant semantic and syntactic information is described in detail in Section 2.1. In Section 2.2, a schematic of the OEM modeling the EOS is introduced, and the details of each stage follow. Olfactory transduction in the AN and Maxillary Palps is modeled with *differential* DNPs in Section 2.3. The glomerular structure of the AL is modeled as multiple processing channels in Section 2.4. 3 *differential* DNPs are described that extract in parallel, respectively, the semantic information and the ON and OFF events that signal the timing of odorant semantics. In Section 2.5, we model the MB Calyx with APL feedback as a *differential* DNP that maps the semantic information into a marked first spike sequence code.

### 2.1 Odorant-Receptor Binding Defines Odorant Identity

As discussed in Section 1, the odorant space largely determines the functional logic of the olfactory circuit. Consequently, we first formally model the semantics and syntax of pure odorants [10].

The proposed odorant space model is not defined by the (largely intractable) chemical structure of the odorants [14, 15]. Rather, it is described by the interaction between odorants and olfactory receptors as a pair of tensors (**b, d**) [8, 9]. As depicted in Figure 2, **b** and **d** denote the 3D tensor of binding and dissociation rates, respectively. In 3D the entries of these tensors are a function of, respectively, the odorant identity, the olfactory receptor type and the index of the OSN that expresses the receptor type. Thus, each entry of **b** represents the binding rate of odorant *o* interacting with olfactory receptor *r* expressed by OSN *n*, where *o ∈* 1, *· · ·*, *O, r ∈* 1, *· · ·*, *R* and *n* = 1, *· · ·*, *N*, and *O* is the the number of odorants, *R* is the number of olfactory receptor types and *N* is the number of OSNs that express a specific receptor type (see Figure 2). **d** is similarly defined.

**Figure 2.**
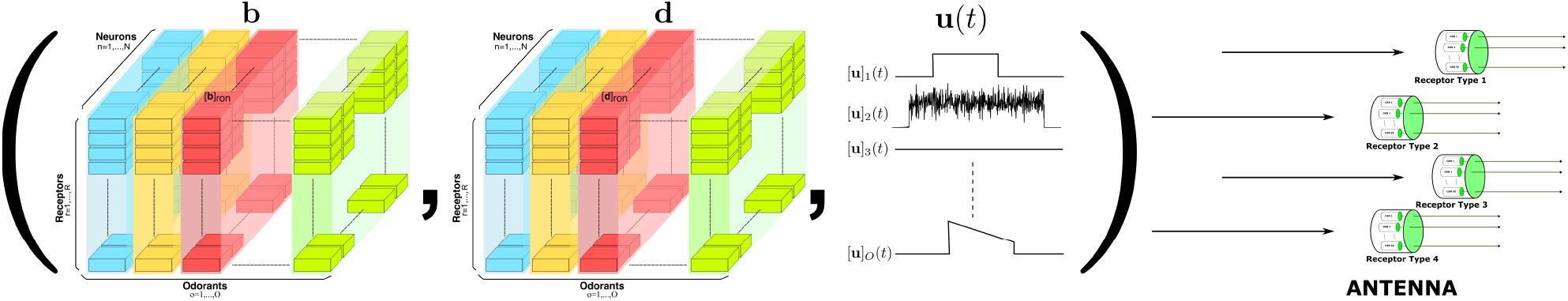
Modeling the space of pure odorants.

The odorant syntax is modeled as a temporal signal, characterized by its odorant concentration waveform *u*(*t*). Even though this signal has no bandwidth limitation as the natural odorant concentration can widely fluctuate in high velocity odorant plumes, the sensilla (see Figure 4(A)) act as a low-pass filter that limits the bandwidth of the concentration waveform [16].

Jointly, the tensor pair determines what types of sensors (olfactory receptors) will be activated by a certain odorant, and the level of activation will be governed by the identity of the odorant in conjunction with its concentration waveform. More precisely, the overall activation of the sensors is determined by the value of the odorant-receptor binding rate modulated by the odorant concentration profile. See Section 2.3 for more details.

In sum, the olfactory objects of the odorant space are explicitly described by both their identity (odorant semantics) and their concentration amplitude (odorant syntax).

### 2.2 Schematics of the Odorant Encoding Machine Modeling the EOS

The circuit architecture of the OEM introduced below consists of a cascade of spatio-temporal *differential* Divisive Normalization Processors (DNPs) [17] each modeling the graph structural pathways of the AN, AL and MB Calyx shown in Figure 1(bottom), and schematically depicted in Figure 3. Note that Figure 1 and Figure 3 taken together represent the 3 workflow modeling steps advanced in [13] consisting of (i) the 3D exploration and visualization of fruit fly brain morphology datasets, (ii) the abstraction of the pathway structure of the connectome and (iii) the creation of canonical, IOPs. The fourth step, the interactive exploration of the functional logic of canonical

**Figure 3.**
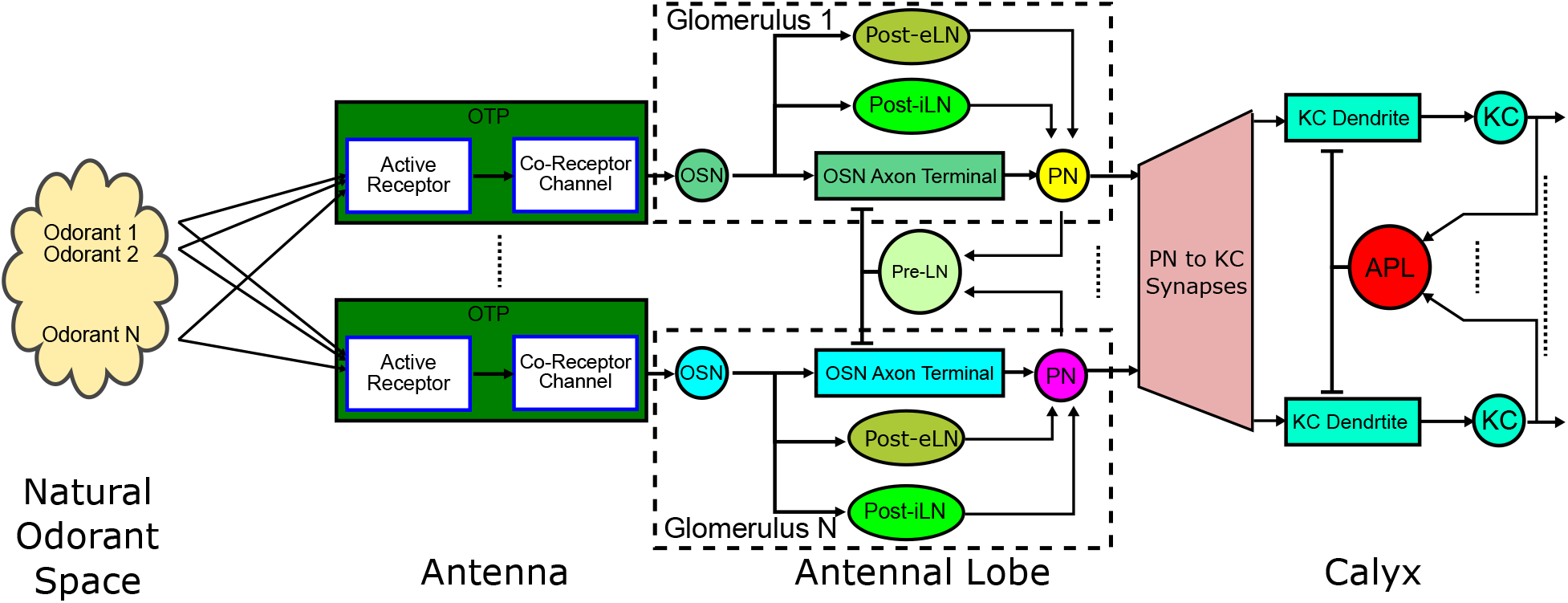
Schematic of the Odorant Encoding Machine (OEM) modeling the early olfactory system of the fruit fly.

*Differential* DNPs are a class of processors that can be used to model sensory processing with a variety of circuit configurations [18, 10]. The simplest *differential* DNPs take the form of a dynamical system

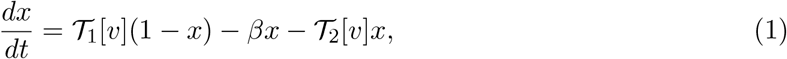

where *v* is the input to the differential DNP, *x* is the output, and *T*_1_ and *T*_2_ are causal Volterra operators [19, 20, 17] of appropriate dimensions. A particular example for *v* appears in Eq. (7). In steady-state, the response amounts to

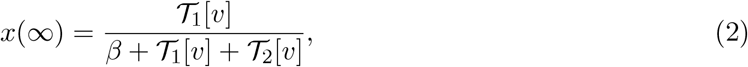

corresponding to a division between two processed version of the input *v* in steady-state. A particular form of Eq. (2) has been considered by [21] as a canonical computation in the brain. Here, *T*_1_ and *T*_2_ can be considered as two processing branches feeding a divisive normalization operator in a feedforward path.

A *divisive* DNP with feedback can be easily obtained by replacing *T*_2_[*v*] with *T*_3_[*x*], i.e.,

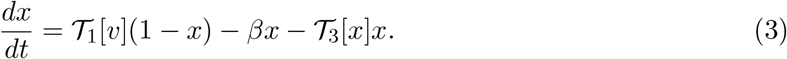

The steady state response if it exists amounts to

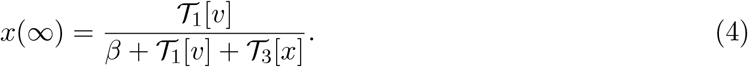

In addition, the feedforward or feedback terms can incorporate global information to construct spatio-temporal *differential* DNPs. For example, by denoting each output with a superscript *i, i* = 1, 2, *· · ·*, *N*, where *N* is the total number of neurons involved in the *differential* DNP, we obtain

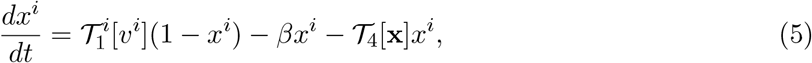

where *T*_4_ receives inputs from all outputs **x** = [*x*^1^, *x*^2^, *· · ·*, *x*^*N*^] ^T^.

Note that in Figure 3, each of the processing stages can be modeled by one or more *differential* DNPs: (i) the Active Receptor and Co-Receptor Channel are modeled with *differential* DNPs with feedback; (ii) 3 types of local neurons in the AL, the Presynaptic Local Neurons (Pre-LNs), the Postsynaptic excitatory LNs (Post-eLNs) and the Postsynaptic inhibitory LNs (Post-iLNs), are modeled as 3 types of *differential* DNPs. (iii) the Calyx features an expansion of PN to KC connectivity, a *differential* DNP modeling a circuit consisting of the KC dendrites, KC biological spike generators and the APL spatio-temporal feedback neuron.

In the rest of this section we describe the above mentioned *differential* DNPs as intelligent olfactory blocks processing in parallel semantic and syntactic information flows.

### 2.3 The Antenna Multiplicatively Couples Odorant Semantics and Syntax

In the first stage of the olfactory transduction process, odorant molecules bind to olfactory receptors expressed by each OSN residing on the Antennae and Maxillary Pulps. The spikes generated by the OSNs are transmitted to the AL. The olfactory transduction, and, the spike generation and transmission run largely in parallel among OSNs. Recent discovery of ephaptic coupling, however, suggests that there are additional lateral interaction between OSNs whose cilia are housed in the same sensillum [22]. Our model currently does not account for interactions between OSNs.

The molecular machinery of odorant transduction is schematically depicted in Figure 4(A). The Olfactory Transduction Process (OTP) model is shown in Figure 4(B).

**Figure 4.**
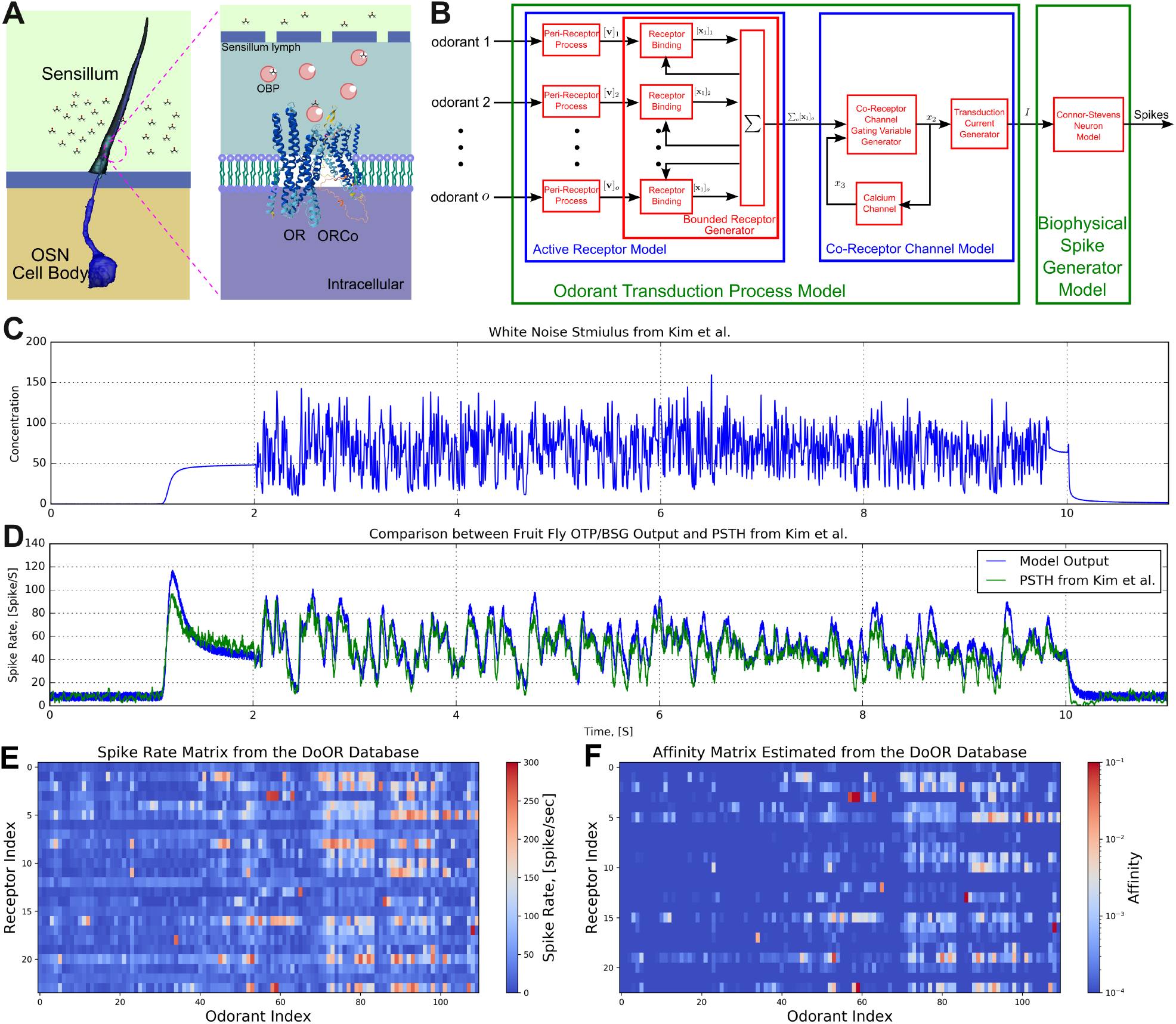
(A) The molecular machinery of odorant transduction. A cartoon of a sensillum that houses OSN cilia. Odorant molecules enter through the sensillum pores into sensillum lymph, to be transported to the membrane of OSN dendrites by odorant binding proteins (OBPs) [23]. The binding of odorant molecules to ORs starts a transduction cascade that opens ion channels located on the OSN cilia. The transduction current then induces the OSNs to generate action potentials. Adapted from [24]. (B) Schematic diagram of the Olfactory Transduction Process for a set *O* of odorant mixture components, *o ∈ O* pure odorant. (C) White noise stimulus waveform [16]. (D) OSN model (blue, [8, 9]) and recorded output (green, [16]). (E) Spike rate matrix from the DoOR database containing 24 odorant receptors and 110 odorants. Each column represents an odorant, and each row represents an OSN receptor type. (F) Each entry of the affinity matrix is estimated from each entry of the spike rate matrix using the inverse of the empirically determined OTP/BSG transformation in steady-state [8, 9]. Note the log-scale color map for the affinity.

The OTP consists of 3 stages. The first stage is the Peri-Receptor Process block modeling the interaction between odorants and the olfactory sensilla before the odorant-receptor binding takes place. The output of this stage can be expressed as

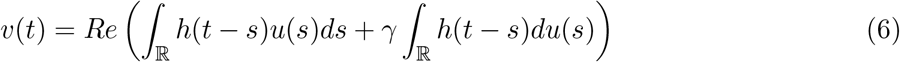

where *Re*(*·*) denotes the rectification function and *h*(*t*) is the impulse response of the peri-receptor process. This process reduces the rapid fluctuations of the odorant concentration waveform and enhances the response to concentration gradients [16].

The second stage shown in Figure 4(B) describes the binding process taking place in the Active Receptor. At any point in time, the number of bound receptors depends on the amplitude of the concentration waveform of the odorant (syntax), and, the binding rate as well as the dissociation rate between the odorant and the olfactory receptors (semantics). This can be mathematically expressed as

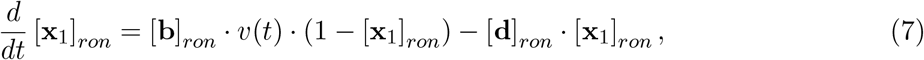

where [*·*]_*ron*_ denotes an entry in the corresponding tensor, and *v*(*t*) is the output of the Peri-Receptor Process block mentioned above. During the binding process, the odorant semantic and syntactic information are multiplicatively coupled and encoded as [8, 9]

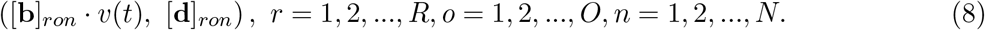

Note that it is not possible to separate the two information streams from the spikes generated by a single OSN, as two different odorants with different concentration waveforms may evoke the same OSN spike response [25]. Thus, without the entangling of the two types of information streams across the population of OSNs, odorants cannot be identified. This is similar to sensing light intensity reflected by an object with a Lambertian surface, which depends on both the luminous intensity and the reflectance of the surface material [26].

The third stage shown in Figure 4(B) is the Co-Receptor Channel that models the opening of transduction ion channels by the total bound receptors. The opening of the ion channels is also controlled by a feedback mediated calcium concentration.

Finally, the transduction current shown at the output of the OTP in Figure 4(B) is encoded by a biophysical spike generator (BSG) into the spike domain.

The OSN model considered here accurately predicts the OSN’s response to the white noise stimulus shown in Figure 4(C). The response is captured in electrophysiology recording shown in Figure 4(D) [16, 8, 9].

Existing OSN recordings in the DoOR database [27, 28] contain responses of 24 odorant receptors to 110 odorants, at a single concentration level. This is depicted in Figure 4(E) where each column represents an odorant, and each row represents an OSN receptor type. This allows to estimate the affinity matrix (the ratio between binding and dissociation rates), show in Figure 4(F), using the inverse of the empirically determined OTP/BSG transformation in steady-state [8, 9]. We note that, only a limited number of recordings are currently available for individually extracting both the binding and dissociation rates. New experiments for identifying the odorant-receptor dissociation rates are in order.

### 2.4 The Antennal Lobe Extracts Semantics and Semantic Timing Events

The Antennal Lobe (AL) consists of some 50 glomeruli. Each glomerulus is the target of OSNs that express the same, olfactory receptor type. Uniglomerular projection neurons (uPNs) also exclusively arborize in only a single glomerulus. For example, in Figure 5(top left), we show the DL3 and VA3 glomeruli. For the DL3 glomerulus, the at4B OSNs (cyan) expressing Or65a project from the AL into the glomerulus, and provide inputs to DL3 PN (magenta), and the synapses are shown in pink. For the VA3 glomerulus, the ab9 OSNs (light green) expressing Or67b synapse onto the VA3 PN (yellow).

**Figure 5.**
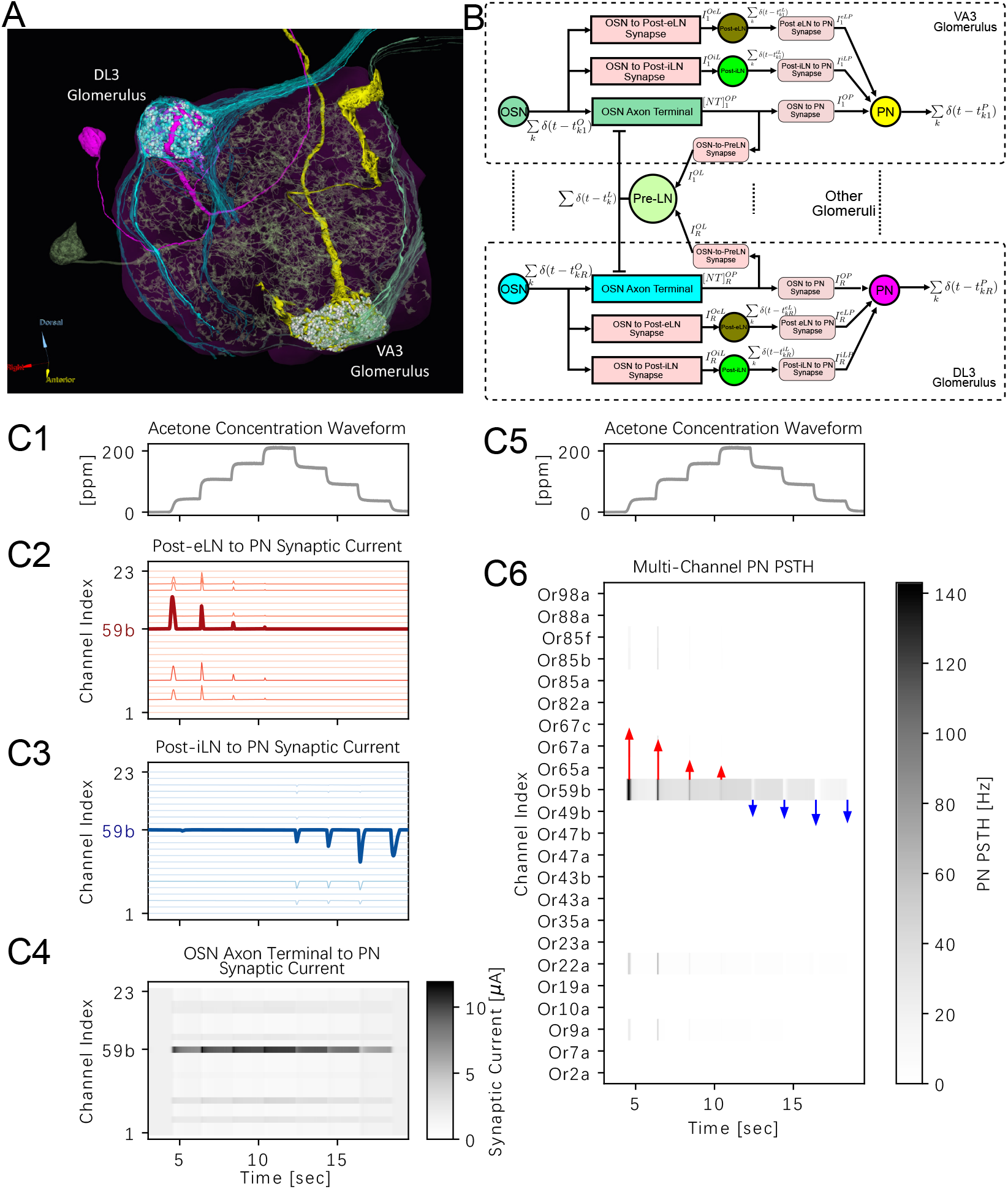
(A) Morphology of a small number of neurons in the AL of the fruit fly brain and (B) a multi-channel AL circuit model. Two glomeruli are shown: DL3 glomerulus with OSNs in cyan and uPN in magenta, VA3 glomerulus with OSNs in light green and uPN in yellow. Synapses are shown in light pink. A Pre-LN is shown in transparent green across the AL. The multi-channel AL circuit model has the same color code for each block. (bottom) an evaluation of the input/output relationship of the AL circuit model has the same color code for each block. (bottom) an evaluation of the input/output relationship of the AL circuit model. (C1,C5) Acetone staircase odorant concentration waveform recorded in [30]. (C2) Post-eLN to PN synaptic currents. Different hues of red indicate strengths of synaptic current, with strongest due to the Post-eLN receiving input from the Or59b OSN indicated in dark red. (C3) Post-iLN to PN synaptic currents. Different hues of blue indicate strengths of synaptic current, with strongest due to the Post-iLN receiving input from the Or59b OSN indicated in dark blue. (C4) OSN Axon-Terminal to PN synaptic current shown as a heatmap. Different hues of grey indicate strengths of synaptic current. Refer to the colorbar for scale. (C6) Multi-channel PN PSTH, with synaptic current of ON/OFF pathways along Or59b/DM4 channel shown as red and blue arrows, respectively. Figure adapted from [10].

In addition to this feedforward connectivity within each glomerulus, a complex network of Local Neurons (LNs) arborize in varying number of glomeruli to facilitate local and global feedback loops within and between glomeruli [29]. In Figure 5(A), we show an instance of a LN (in transparent light green) that innervates all glomeruli. In [29], we have characterized the interaction of LNs with OSNs and PNs in each glomerulus, showing a diverse range of patterns. Some LNs receive feedforward inputs from OSNs and feedback inputs from PNs while providing feedback to both. Others may have a subset of these connections. While the PNs receive dedicated inputs only from a single type of OSNs, LNs enable crosstalk between them. Thus, the AL outputs carried by PNs are the result of inter-glomerular processing.

The LN connectivity in the AL appears to be too complex to be used directly in revealing their individual functions. Going the extra mile, our first step is to postulate the overall functionality of the different classes of LNs based on some of the connectivity features observed in the AL connectome.

Building on the 2D encoding model of OSN transduction [9], *differential* DNPs have been explicitly designed to extract odorant semantic and syntactic information from olfactory signals carried by OSNs [10]. We advanced temporal *differential* DNPs for single-channel AL circuits and spatio-temporal *differential* DNPs for multi-channel AL circuits, incorporating three types of local neurons that shape the information flow within the AL: (i) pre-synaptic pan-glomerular inhibitory local neurons (Pre-LNs), (ii) post-synaptic uni-glomerular excitatory local neurons (Post-eLNs), and (iii) post-synaptic uni-glomerular inhibitory local neurons (Post-iLNs).

For single-channel AL circuits modeled as 3 parallel temporal *differential* DNPs (see Figure 5(B) within each glomerulus dashed box), we found that the divisive normalization mediated by the Pre-LN plays a central role in enhancing the *concentration-invariance of the PN responses*, which we interpret as a first step toward extracting odorant semantics. In parallel, Post-eLN excitation and Post-iLN inhibition selectively enhance transient PN responses at stimulus onset (ON) and offset (OFF), thus extracting the ON-OFF timing of the syntactic information.

Scaling from single-channel to multi-channel AL circuits (see Figure 5(B)), concentration-invariant PN population responses can be achieved across all channels. We hypothesized that this invariant PN activity corresponds to a reconstruction of the odorant semantics defined by the odorant *affinity rate* vector. Each entry of the affinity rate vector is the element-wise ratio between the odorant-receptor binding and dissociation rates defined in Section 2.1. The contrast-boosted transient components of PN responses encode ON and OFF timing events associated with the presence of odorant semantics (identity) across all channels (see Figure 5(C2,C3)). Note that the spatio-temporal *differential* DNP along the Pre-LN pathway is critical for the recovery of the odorant semantics (identity) at the level of PNs (see Figure 5(C4)). Together, the multi-channel AL circuits disentangle the confounded semantic and syntactic information streams at the output of the AN, and encode both the identity and the timing information of the odorant objects into the PN output spike trains (see Figure 5(C6)). The resulting PNs spike trains carry both the “what” (identity) and the “when” (timing) about the odorant semantics. Both of them are the key ingredients of associative learning in the MB.

In summary, the functional logic of the AL is that of an *ON-OFF odorant object identity recovery* processor, robustly extracting the temporally marked odorant semantics [10].

### 2.5 The MB Calyx Maps Odorant Semantics into a Marked First Spike Sequence Code

The MB Calyx largely consists of the axons of the uniglomerular PNs, dendrites of the Kenyon Cells and the multi-input multi-output APL neuron (see Figure 6(A)). PN axons terminals (Figure 6(A) in yellow, blue and magenta) form large boutons in the Calyx, with numerous presynaptic sites providing inputs to many KCs and the APL neuron. PN boutons also receive feedback from the APL neuron. Synapses around a PN bouton form what is collectively called a microglomerulus [31]. Each of the KCs, on the other hand, forms several discrete dendritic claws (Figure 6(A) in turquoise). Each claw is typically associated with a single PN bouton. On average, each KC has 6 to 7 claws, connecting to the same number of different PNs. Thus, with a total of ~2,000 KCs and ~50 types of uniglomerular PNs, the olfactory signal generated by the PNs undergoes an expansive transformation.

**Figure 6.**
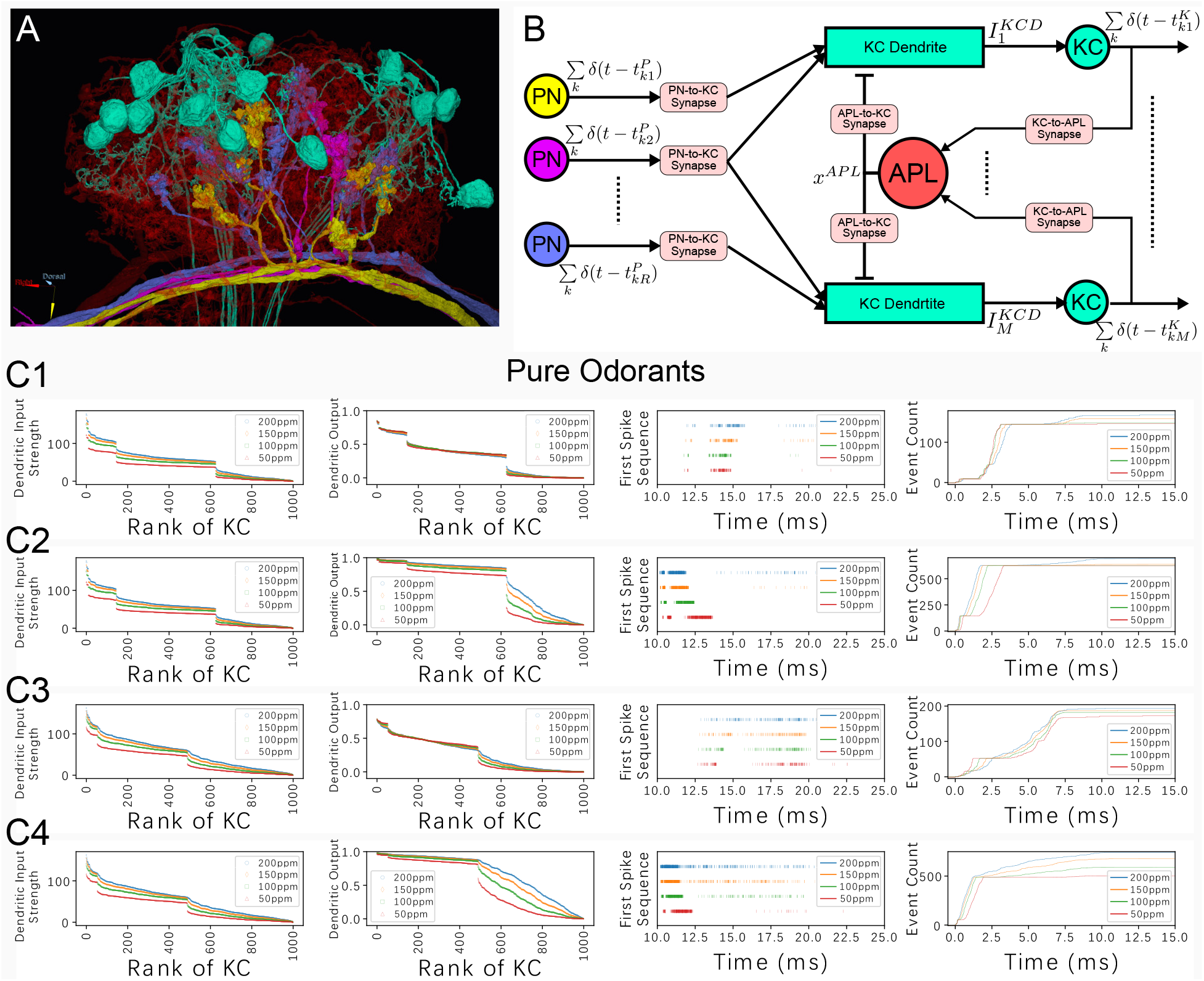
(A) Schematic morphology of a small subset of neurons in the MB Calyx. (red) APL, (turquoise) KCs, (other colors) PNs. (B) Schematic diagram of the MB Calyx circuit with spatio-temporal APL feedback. (C1-C4) Response in the MB Calyx circuit to different pure odorants. (C1) Furfural, (C2) Furfural with APL feedback silenced, (C3) 1-Butanol, (C4) 1-Butanol with APL feedback silenced. For each pure odorant, 4 different constant concentrations are included: 200ppm, 150ppm, 100ppm and 50 ppm. **(1st column)** Ranking of KC dendritic inputs. **(2nd column)** Ranking of KC dendritic outputs. **(3rd column)** Odorant semantics encoded in the time domain across the population of KCs. The first spikes of each of the active KCs in response to each odorant are collected onto a single row for each of the odorant concentration amplitude values. **(4th column)** Cumulative inter-spike interval of the first spike sequence

In addition to PN boutons and KC dendrites, an APL neuron (Figure 6(A) in red) covers the entire MB, including the Calyx. It receives inputs from almost all PN boutons and all KCs, and also provides them with feedback.

We map the MB Calyx circuit into a circuit model, whose schematic diagram is shown in Figure 6(B). Note that the color code in the circuit diagram is consistent with the color code in the visualization of the morphology of the connectome. For simplicity, the PN inputs are linearly summed [32]. We notice that the summed PN inputs are still concentration dependent, as shown in Figure 6(C1-C4, column 1).

We model the APL to KCs feedback circuit as a *differential* DNP, as shown in Figure 6(B). This DNP largely reduces the concentration dependency at the KC dendritic output, demonstrated in Figure 6(C1 and C3, column 2). In contrast, by silencing the APL neuron and thereby removing the feedback of the MB Calyx circuit, the KC dendritic output no longer exhibits sparseness. Neither is it distinct across different odorants (see Figure 6(C2 and C4)). Additional DNPs modeling the interaction between PN boutons and the APL neuron can similarly be constructed. We omit them here for simplicity.

With the feedback driven by the APL neuron included, we simply merge the first spike of each of the KC responses to form a *marked first spike sequence code*, as shown in Figure 6(C, column 3). Each spike is marked by the KC that generates it. This code reflects the ordered strength of the concentration-invariant KCs dendritic output. Codes such as the primacy code utilizing ordered strength of a population of neurons have also been proposed for the outputs of the AN and AL [33, 34, 35, 36, 37], as well as in the piriform cortex of vertebrates [38], and proposed for modeling circuits in the visual system [39, 40]. Note that the cumulative inter-spike interval of the marked first spike sequence code, shown in Figure 6(C1 and C3, column 4), is highly conserved for different concentrations of the same odorant, but distinct for different odorants. This diversity, however, is largely reduced by the absence of the APL neuron feedback (see Figure 6(C2 and C4, column 4)). See Supplementary Figure S1 for an extensive comparison (for all tested odorants) of the MB Calyx circuit with and without the APL neuron feedback.

The expansion of the connectivity from the PNs to KCs and the readout of the input dendritic ordered strength is a simple yet effective strategy for extracting features of the AL output vector of the PNs. Moreover, the readout of the KC code by the downstream MBONs is similarly kept simple. MBONs receive inputs from all KCs that project to the corresponding compartment. The code can generally be applied to different flies even though the connection between PNs and KCs is random and differs among flies.

## 3 Classification of Pure Odorants

The output of the OEM schematically depicted in Figure 3 feeds into the MB associative memory circuit. The MB circuit must be able to, from the marked first spike sequence code, distinguish different odorants in the form of a memory hit or the absence of it.

In this section, we advance a rank-based distance measure to evaluate how the new representation, the marked first spike sequence code, can be used in distinguishing ON-OFF odorant semantics.

### 3.1 Rank-Based Distance Measures for ON-OFF Odorant Semantics

The outputs of the KCs feed directly into the associative memory circuits of the MB compartments. There, the ON-OFF odorant semantics carried by the KCs and the sensory semantics carried by the Dopaminergic neurons (DANs) are integrated into the associative learning process. Olfaction is a highly effective sensory modality that helps the fly associate ON-OFF olfactory semantics with non-olfactory semantics, such as an electric shock and a taste of sugar. Furthermore, sensory semantics such as starvation or cold can also be recalled by the ON-OFF odorant semantics alone.

As schematically depicted in Fig. 7, the MB distinguishes currently experienced ON-OFF odorant semantics from the stored semantics [41]. In other words, the MB recalls sensory semantics from previously associated odorant semantics regardless of its syntax thereby achieving separation between different odorant identities. We refer to this as semantic specificity.

**Figure 7.**
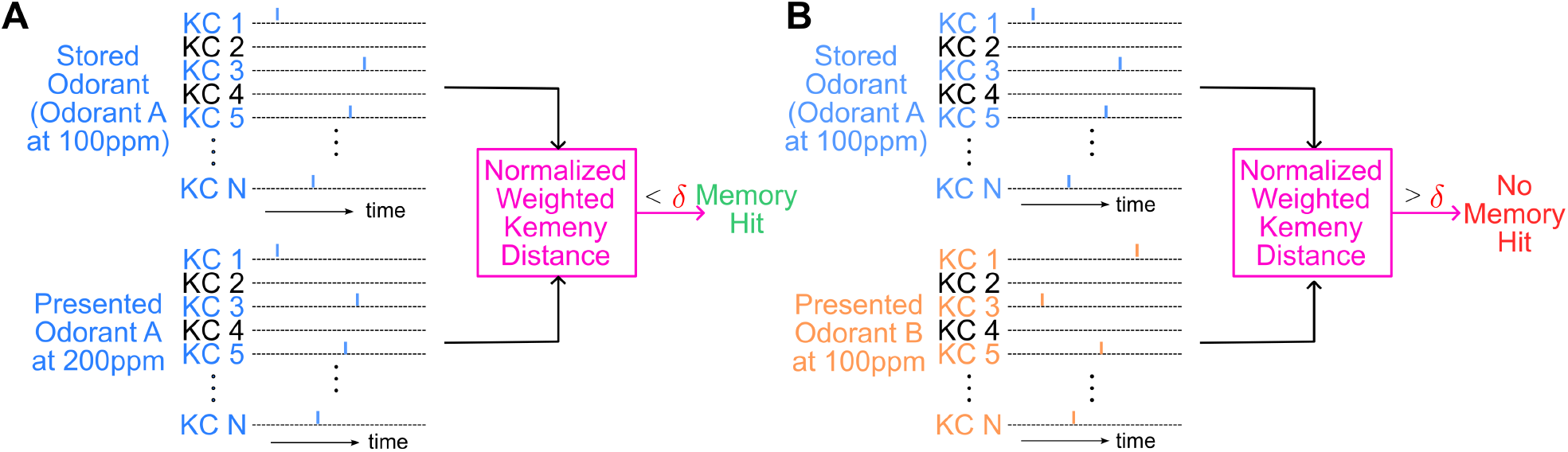
(A) top: The marked first spike sequence code of the odorant that has already been previously stored (associated) at 100ppm concentration. bottom: The marked first spike sequence code of the same odorant at 200pm should, in spite of the concentration difference, result in a low distance measure between the two spike sequence codes, leading to a memory hit. (B) top: Same as top of (A). bottom: A different odorant will result in a greater distance measure to the stored odorant semantics, signaling that the odorant presented has not been stored.

Here, we incorporate KC labels as marks into the first spike sequence code. More precisely, we mark each spike in the first spike sequence code with the label of the KC, *i*.*e*., the *i*-th spike is marked as (*t*^*i*^, *r*^*−*1^(*i*), where *r*: *S →* ℕ is the rank function that maps each KC in the set of all KCs to a rank (assuming that there is no tie and thus *r* is invertible). The marks provide the ranking information of the KC neurons that generated the first spikes in the sequence.

By incorporating the rank information, we can now employ a rank-based distance metric, such as Kemeny-Snell [42] or Emond-Mason [43], between two marked first spike sequence codes in order to quantify their difference to be used for matching a just sensed odorant with the one stored during associative learning.

Consider a set of *m KCs* whose times to first spike in response to odorant *A* is listed as the set 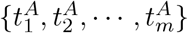 where 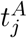 is the first spike time of the *j*-th KC. We construct the (*m, m*) score

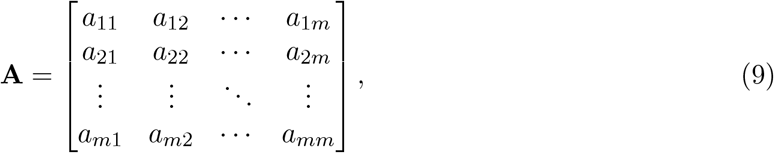

Where

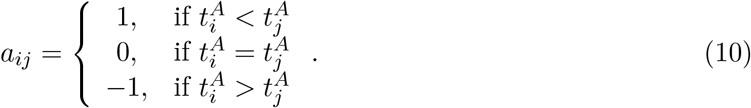

Similarly, we define the matrix **B** for the first spike times 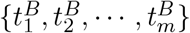 of *m* KCs responding to odorant *B*. The Kemeny-Snell distance between the two sequences of first spike times is given by

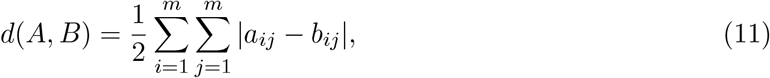

and the rank correlation coefficient *τ* amounts to [43]

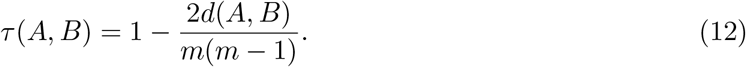

Note that Kemeny-Snell distance *d* and rank correlation coefficient *τ* have been used primarily in studying rank aggregation to find a consensus ranking from a number of preferential rankings. To increase the sway of KCs that spike earlier, we use a weighted rank correlation measure [44], a variation of the Kemeny-Snell distance measure. We refer to this metric as the *normalized weighted Kemeny distance* (NWKD). We use the NWKD to define the distance between two marked first spike sequence codes. See the Appendix for a detailed mathematical treatment.

### 3.2 Classifying Marked First Spike Sequence Codes with the NWKD Metric

We ask if the identity of an odorant (semantics) represented by the marks of the first spike sequence code, when stored in the MB associative memory, can be recalled by the same odorant but not others. As an example, in Figure 8(A1), we consider acetone as the associated odorant and compute the distance between the marks of the marked first spike sequence code of acetone and the marks of all the other 103 odorants. The proposed rank-based distance between two odorants is calculated for the same odorant pair at different concentration levels. The minimum NWKD for each pair across all concentration levels defines the distance between two odorants. The histogram of the evaluated distances is shown in Figure 8(A1) on the top in blue, as the count of the number of odorants within a bounded distance (0.02 on the x-axis) to acetone.

**Figure 8.**
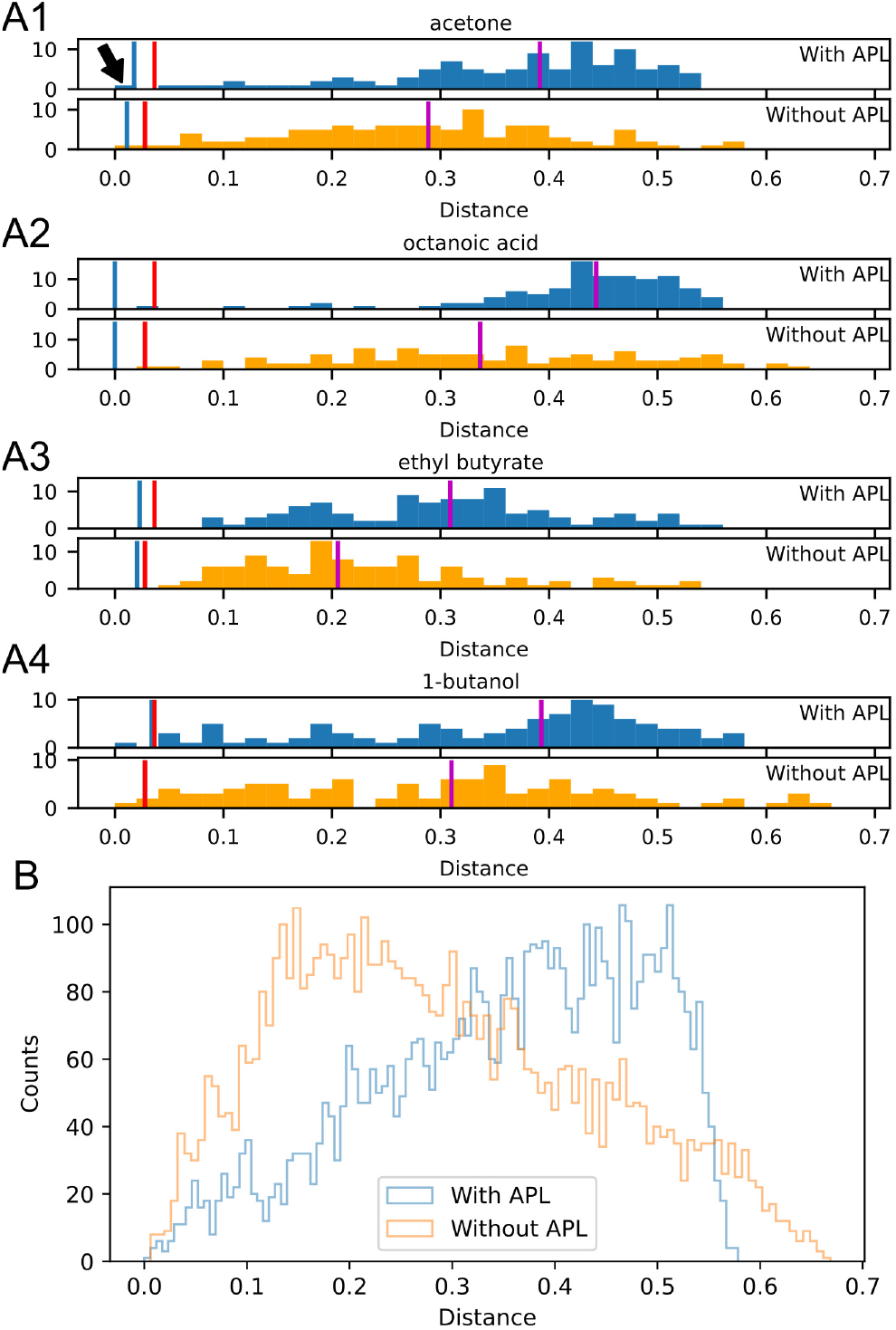
Role of APL feedback in shaping distances between the first spike sequence codes. (A1) Histograms of distances between the marked first spike sequence code of acetone and the other 103 odorants. The y-axis is in the unit of counts of the number of odorants. Blue vertical line indicates the maximum distance between first spike sequence codes responding to acetone at different concentration levels. (Top) KC responses of a MB Calyx model with APL feedback. (Bottom) KC responses of a MB Calyx model without APL feedback. Red vertical line indicates the maximum value of all blue lines across all odorants, including those in A2-A4 and in the Supplementary Figure S2. Purple lines indicate the median value of the NWKD between the odorant and all other odorants. (A2-A4) Same as A1 but for octanoic acid, ethyl butyrate and 1-butanol, respectively. (B) Histogram of distance measures between the marked first spike sequence codes of all pairs of odorants. Distances calculated from KC responses (blue) of a MB Calyx model with APL feedback and (orange) without APL feedback.

We also computed the maximum NWKD between the marked first spike sequence codes from responses to acetone pairs of different concentrations. This is indicated by the blue vertical line. Any odorant whose distance to acetone is to the left of the vertical line cannot be distinguished from acetone.

A histogram is also shown for 1-butanol (Figure 8(A2)), ethyl butyrate (Figure 8(A3)) and octanoic acid (Figure 8(A4)). Again, the blue vertical lines indicates the maximum NWKD across different concentrations of that same odorant. That is the distance under which another odorant cannot be distinguished from the named odorant. The histogram for all odorants is shown in Supplementary Figure S2. We obtained the red vertical line by taking the maximum of the blue vertical lines across all odorants listed. The red vertical line is a global distance threshold. Any odorant whose distance to a stored odorant is above the red vertical line can be safely considered as not to have been associated.

Two cases are considered in Figure 8(A1-A4): distance histograms for marked first spike sequence codes obtained with/without APL feedback. On top of each panel (with APL) we can see that, the stored odorant, *e*.*g*., acetone for panel A1, can be largely differentiated from the other odorants by the proposed distance measure, except for 2-butanone indicated by the arrow, whose affinity profile is largely the same as the one of acetone for the 23 odorant receptors model we used here [27]. This exception may be dropped when the full spectrum of 50 odorant receptors is available. When the APL neuron is silenced, the rank distances produced by the circuit with APL feedback present (shown in the histogram at the bottom of each panel A1-A4 in Figure 8) are closer to 0.5, indicating that the APL neuron decreases the correlation between the marked first spike sequence codes of different pure odorants.

This trend can be observed for almost all odorants (see Supplementary Figure S2). Figure 8(B) shows the distance histograms between all 5,356 pairs of the 104 odorants tested. We can see a clear shift of the histogram of pairwise distances when APL feedback is present in the MB Calyx circuit. The APL feedback visibly shifts the distance between the marked first spike sequence codes for different odorants toward 0.5, a value that indicates no correlation (see also Appendix A), the classification error will decrease. This suggests that the circuit with APL feedback potentially leads to a better classification of the marked first spike sequence codes, thereby enhancing the underlying odorant semantic specificity.

The result shown here suggests that the marked first spike sequence code is a strong candidate for the representation of odorant semantics at the output of the MB Calyx. Across all odorants shown in Supplementary Figure S2, only 65 out of 10712 calculated distances are to the left of the global threshold (red vertical line). In other words, the error rate of mistaking a different odorant as the stored odorant is only 0.61%. Thanks to the APL feedback, this error rate can still be kept low when more odorants are considered. Compared to a binary code with activated KCs labeled as 1 and 0 otherwise [45, 46], the rank-based distance measure can further differentiate odorants as different ranking can be placed on the same set of activated KCs. Yet, the marked first spike sequence code can naturally carry such rank information without further effort required for extracting the ranking.

In Figure 6, we have argued that the feedback from the APL neuron is capable to largely normalize the range of KC outputs across different concentrations, leading to a more “standard” output from a wide range of inputs. To a degree, the evaluation of the role played by Pre-LN feedback in the AL (see Figure 5) is similar as both feedback circuits are *extracting* ON-OFF odorant semantics.

Taking it one step further, the classification of the marked KC first spike sequence codes provides a novel way of evaluating the effectiveness of the (large scale MIMO) APL feedback neuron in the MB Calyx circuit. The rank-based distance measure allows for the *comparison* between the extracted semantics, a key aspect of olfactory intelligence. The effectiveness of the APL feedback neuron in shaping distances between the first spike sequence codes highlights the key role played by the the EOS in making the natural world of odorants intelligible.

## 4 Discussion

The premise that time carries information in the nervous system was placed on firm experimental footing thirty years ago by [47], who showed that cortical neurons driven by fluctuating currents emit spike trains reproducible to better than a millisecond, whereas the same neurons driven by constant current fire with higher variance. Those experiments, however, necessarily interrogated one neuron at a time. What they established is that the single neuron is a faithful channel; what the precisely timed spikes are *about* is a question that a single neuron, in isolation, cannot answer. Making the world of odorants intelligible calls for more than a reliable channel; it calls for a circuit.

It is the pathway from the AN through the AL to the MB Calyx, and the feedback loops embedded within it, that turns the odorant waveform into a temporal code whose ordering carries odorant semantics. The timing of a single KC spike acquires meaning only when read against the timing of the population to which it belongs. Thirty years on, the question has thus shifted from whether spike timing can carry information to what a circuit must compute for spike timing to render the world intelligible.

To investigate how spike timing renders the natural world of odorants intelligible, we explicitly modeled odorant stimuli by theoretically and computationally characterizing their object identity (“semantic information” or “semantics”), and concentration waveform (“syntactic information” or “syntax”) [8, 9, 10]. Under this model, semantic information is characterized by a time-independent spatial tensor, and therefore, odorant object identity is encoded collectively with spatially-distributed sensors. Odorant syntax, on the other hand, is time-varying and embedded in the individual sensor responses. Elements of olfactory intelligence arise already in the processing of these two streams of information by the IOPs through the early olfactory stages.

A key first element of olfactory intelligence in the *Drosophila* is the binding of odorant objects to receptors expressed in the OSNs. The odorant semantics and syntax are multiplicatively coupled during the process of olfactory transduction by the OSNs in the AN. This forms a baseline for the representation of odorant semantic and syntactic information at the first 3 stages of early olfactory transduction and processing: the AN, AL and MB Calyx circuits.

The IOPs modeling the OTP in the AN consist of the active receptor and the co-receptor channel blocks. The spike generation and transmission processes corresponding to multiplicatively coupled odorant semantic and syntactic information run largely in parallel among OSNs.

The IOPs employed in the AL act upon the confounding representation of the odorant semantic and syntactic information at the output of the AN. In parallel, the IOP modeling the Pre-LN feedback circuit extracts the semantic information - the identity of the odorant object, and the one modeling the Post-eLNs and Post-iLNs extracts the ON-OFF timing events to signal when an odorant object emerges and when it fades away. Timing of this processing parallelism raises a number of questions regarding the processing in the AL.

The IOP model of the MB Calyx consisting of the PN-KC expansion circuit and APL feedback circuit provides a novel representation of the odorant semantics as a marked first spike sequence code. The code, starting at the ON timing event and through the time interval [ON, OFF], provides odorant semantic information to the MB. This is precisely what an accurate spike timing at signal onset can support [47]. With the novel rank-based distance measure we advanced here, the odorant semantics represented by the marked first spike sequence code can be effectively classified to support memory operation in the MB and allows a low complexity rapid readout.

Overall, our study of the early olfactory pathway is anchored in time and abides by causality. Time underlies the definition of the odorant syntax, the processing with *differential* DNP circuit, and is the unit used by the representation of the odorant semantics in the marked first spike sequence code. Without considering a causal odorant processing pathway, models of the olfactory system are incomplete for uncovering the functional logic of the early olfactory circuits.

Furthermore, the natural representation of semantics accessing the MB associative memory circuits lies in the spike domain. Time is an intrinsic variable of the concentration waveform that is critical during associative learning [48]. Further characterization of the role played by the marked first spike sequence code in forming associative memory is in order.

We modeled these causal circuits with *differential* DNPs, described by non-linear differential equations with largely stable temporal and spatio-temporal feedback loops. The resulting cascade of feedback circuits in the EOS are instrumental in separating odorant semantics from the confounding ON-OFF odorant syntactic information. Feedback circuits are extremely common in the fruit fly brain across modalities [18, 49], and those built upon large scale MIMO neurons, such as the APL neuron, call for novel methods of end-to-end evaluation/classification. By advancing one evaluation/classification method here based on the information carried by the marked first spike sequence code at the output of the MB Calyx circuit, we opened up a new research direction in feedback control/processing of semantic information.

Another set of important questions in olfactory intelligence arises in dealing with odorant mixtures. As previously observed in many olfactory systems, odorant mixtures can be perceived as configural or elemental [50, 51, 52], and receptor binding of odorant mixtures is more complex [53, 54, 55]. In addition, olfactory processing already starts with ephaptic coupling between the OSN axons [22, 56, 57]. This has important implications on the processing of odorant semantics. Under what conditions would an odorant mixture be considered as a new odorant, and when would it be possible to identify the odorants composing a mixture? These questions apply when the mixture is perceived as a single odorant object. An odorant mixture also arises when multiple odorant objects are presented at the same time [58]. This is similar to the cocktail party problem in audition, as the task of the olfactory system is to recognize a particular odorant or a mixture from a background of odorant mixtures. How can odorant semantics be defined and how does odorant syntax play a role in recognizing these separate odorant objects? Under what conditions can an odorant be overshadowed by a mixture of odorants? The answer may lie in a more in-depth modeling of the space odorant mixture semantics and syntatics as well as the feedback circuits that operate on ON-OFF semantic information in higher neuropils of the fruit fly brain.

To conclude, the EOS is not merely a front end that relays odor information to the seat of associative memory. Olfactory intelligence is already at work in the first three stages of the EOS, each contributing distinct elements to making the natural world of odorants intelligible.

## Abbreviations

(PNs): Project Neurons
(KCs): Kenyon Cells
(APL): Anterior Paired Lateral
(MBONs): Neuron, Mushroom Body Output Neurons

## Acknowledgments

The results reported here were presented in part at the *Workshop on the Functional Logic of Neural Circuits: Diamonds in the Rough*, San Juan, Puerto Rico, February 28, 2024, at the *Computational Neuroscience Meeting*, CNS*24, July 20-24, 2024, Natal, Brazil, at the *Symposium on Informers: Computational and Organizational Insights from the Insect Nervous System*, Entomology 2024, Phoenix, AZ, November 10-13, 2024, and at the *Workshop on the Nature of Intelligence, Bridging Animal and Artificial Intelligence*, 4–5 September 2025, University of Sheffield, UK.

The research reported here was supported, in part, by the National Science Foundation under grant #2024607 and grant #2400687.

## A A Brief Definition of the Distance Measure of the Marked First Spike Sequence Codes

In this section, we provide a brief introduction to the rank-based distance measures. We have introduced the Kemeny-Snell distance measure [42] in Eq. (11) and the rank correlation coefficient *τ* in Eq. (12) to provide additional intuition behind the metric.

The basic intuition behind the rank-based distance measure is that, a rank swap between a pair of adjacently ranked KCs will result in a distance of 1. For example, for 4 KCs *A, B, C* and *D*, the ranking order *ABCD* differs from *ACBD* or *ABDC* by the same Kemeny-Snell distance, because there is exactly one swap between a pair of adjacently ranked items in each case. The overall distance is the minimum number of such swaps needed to transform one rank order into another.

However, since the KCs that fire earlier receive inputs from more active, and supposedly more “important” PNs, swapping the ranking between these KCs should have higher impact on the distance than swapping two KCs that fire later. For example, the distance between *ABCD* and *ACBD* should be greater than that between *ABCD* and *ABDC*.

To achieve this, we now extend the distance measure between two marked first spike sequence codes to the weighted rank correlation coefficients [44].

We first note that *τ* in Eq. (12) can also be computed in a more tractable way (for optimization purposes) as

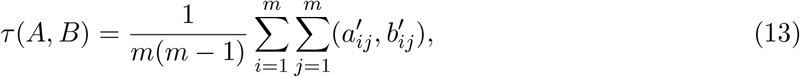

where the score matrix is constructed with elements

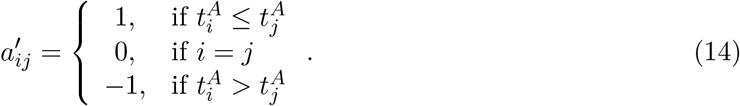

Let *π* be a permutation of the index set (1, 2, …, *m*). By applying *π* on the KC index set, we obtain 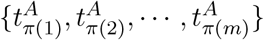. We now construct a pair of matrices with a permutation *π* in response to odorant *A* as

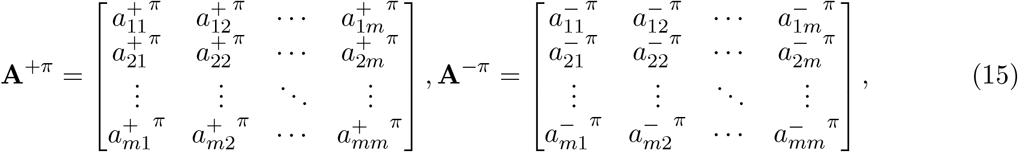

where

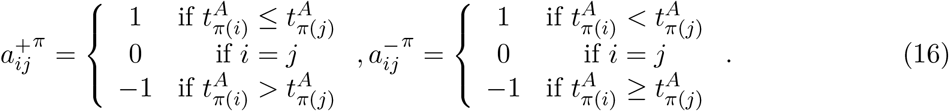

Note that the only difference between 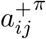 and 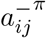 is whether 1 or *−*1, respectively, is assigned to the tied spike times.

We now define the permutation *π*_*A*_ that maps the KC index set into the non-decreasing first spike time sequence 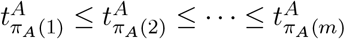. We denote the set of first spike time KC responses to odorant *B* as 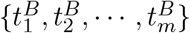. The permutation *π*_*B*_ maps the KC index set into the non-decreasing first spike time 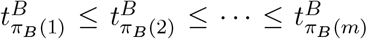. The weighted correlation coefficient between the first spike sequences in response to odorants *A* and *B* is defined, by abuse of notation, as

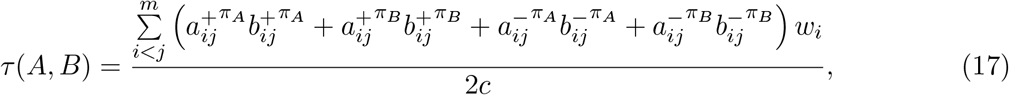

where **w** = [*w*_*1*_, *w*_*2*_, *w*_*m−*1_] is a non-increasing weighting vector, and *w* is the weight given to position *i* in the ranking, and, 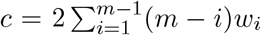 is the maximum weighted Kemeny distance [59].

Again, by abuse of notation, the normalized distance between the KC responses to odorants *A* and *B* amounts to

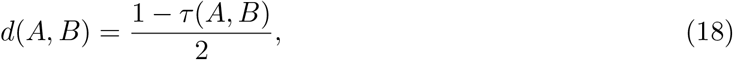

and takes values in [0, 1]. *d*(*A, B*) = 0 indicates that the two first spike sequence ordering are the same. *d*(*A, B*) = 1 indicates the exact reverse order between the responses to *A* and *B. d*(*A, B*) = 0.5 will result in a correlation of 0, thus making the responses to *A* and *B* uncorrelated.

In Section 3, we chose *w*_*i*_ = 0.95^*i−*1^, *i* = 1, 2, *· · ·*, *m −* 1, and further normalized the weights with the constraint 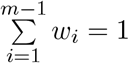.

