## Supplementary Figures for "Elements of Olfactory Intelligence in *Drosophila*"

### Elements of Olfactory Intelligence in *Drosophila*
### Supplementary Figures

Aurel A. Lazar[1†*] and Yiyin Zhou[2†]

[1] *Department of Electrical Engineering, Columbia University, New York, NY 10027, USA*
[2] *Department of Computer and Information Science, Fordham University, New York, NY 10023, USA*
[†] *Authors' names are listed in alphabetical order.*

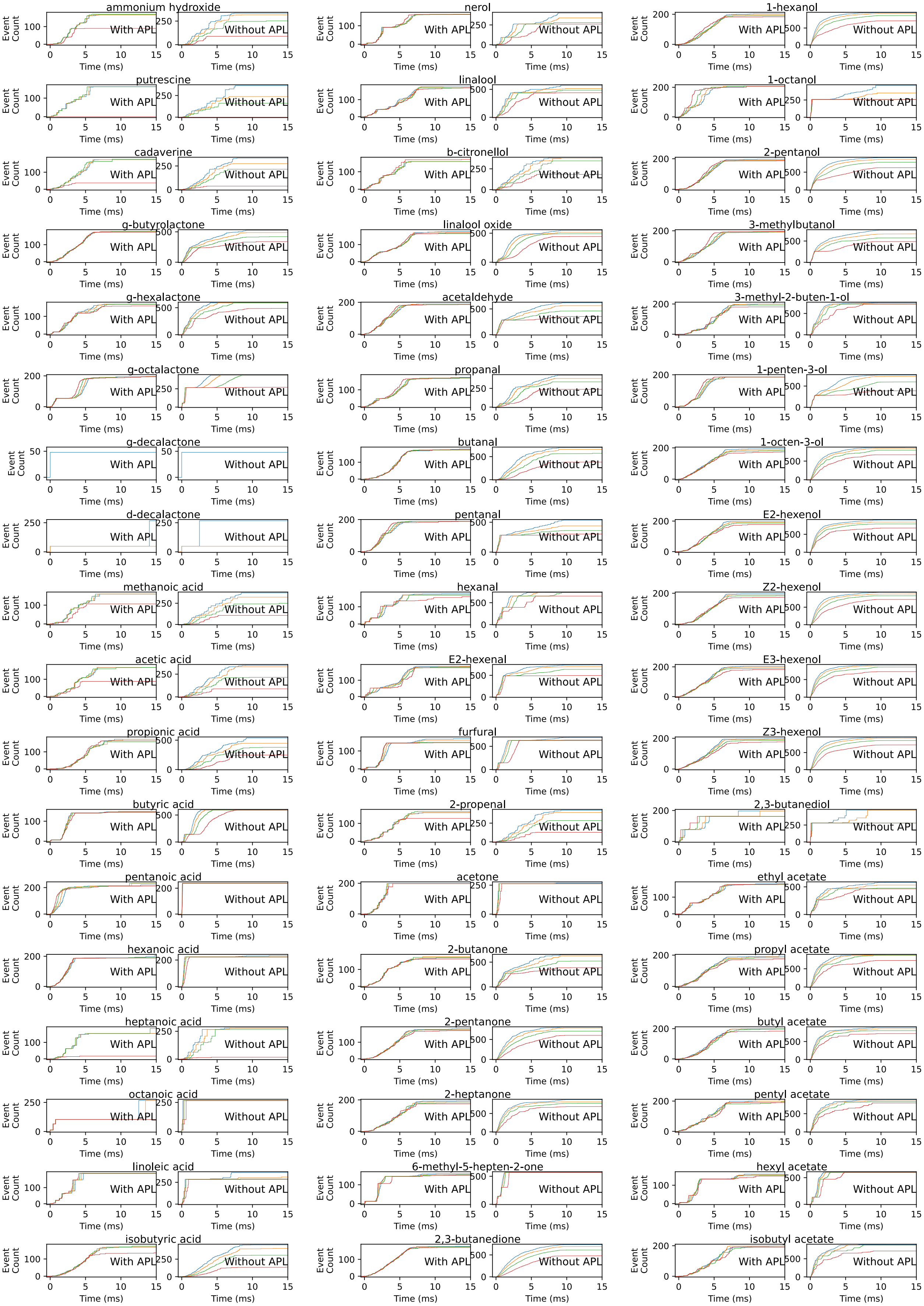
ammonium hydroxide
putrescine
cadaverine
g-butyrolactone
g-hexalactone
g-octalactone
g-decalactone
d-decalactone
methanoic acid
acetic acid
propionic acid
butyric acid
pentanoic acid
hexanoic acid
heptanoic acid
octanoic acid
linoleic acid
isobutyric acid
nerol
linalool
b-citronellol
linalool oxide
acetaldehyde
propanal
butanal
pentanal
hexanal
E2-hexenal
furfural
2-propenal
acetone
2-butanone
2-pentanone
2-heptanone
6-methyl-5-hepten-2-one
2,3-butanedione
1-hexanol
1-octanol
2-pentanol
3-methylbutanol
3-methyl-2-buten-1-ol
1-penten-3-ol
1-octen-3-ol
E2-hexenol
Z2-hexenol
E3-hexenol
Z3-hexenol
2,3-butanediol
ethyl acetate
propyl acetate
butyl acetate
pentyl acetate
hexyl acetate
isobutyl acetate
With APL
Without APL
Event Count
Time (ms)

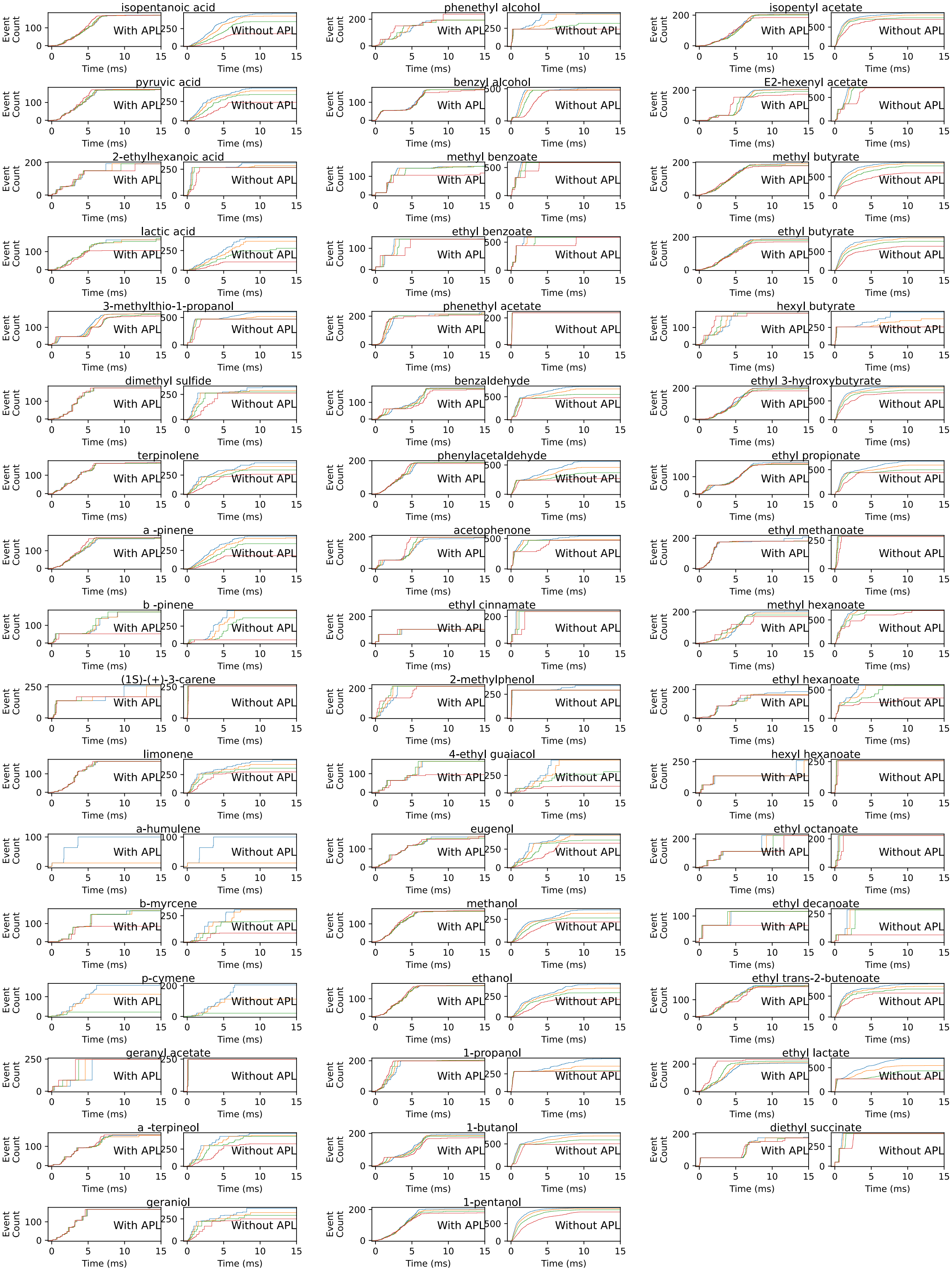


**Figure S1:** Comparison between cumulative inter-spike interval statistics of the first spike sequence code generated by the Calyx circuit with (left in each panel) and without (right in each panel) APL feedback neuron for each odorant.

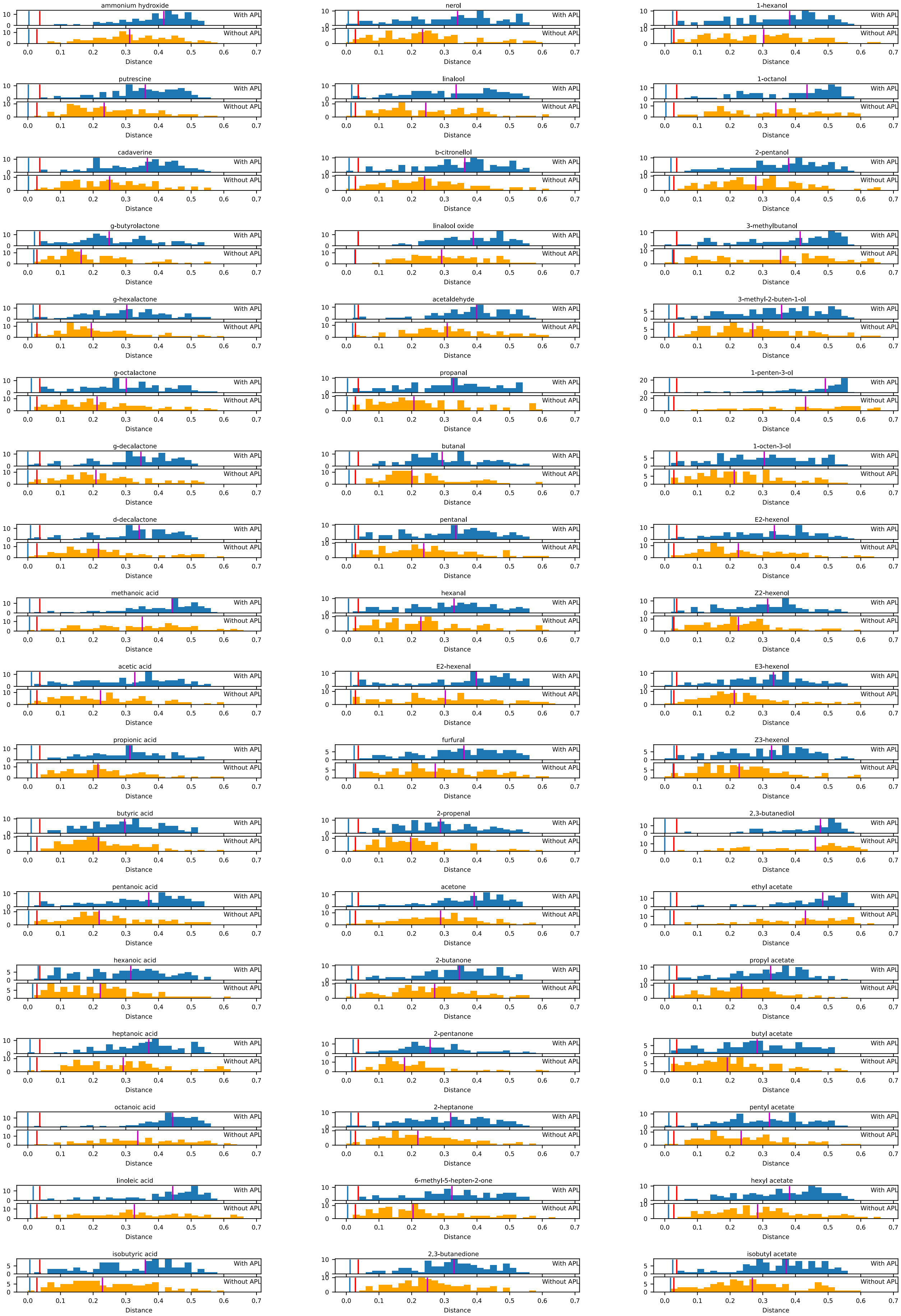
ammonium hydroxide
putrescine
cadaverine
g-butyrolactone
g-hexalactone
g-octalactone
g-decalactone
d-decalactone
methanoic acid
acetic acid
propionic acid
butyric acid
pentanoic acid
hexanoic acid
heptanoic acid
octanoic acid
linoleic acid
isobutyric acid
nerol
linalool
b-citronellol
linalool oxide
acetaldehyde
propanal
butanal
pentanal
hexanal
E2-hexenal
furfural
2-propenal
acetone
2-butanone
2-pentanone
2-heptanone
6-methyl-5-hepten-2-one
2,3-butanedione
1-hexanol
1-octanol
2-pentanol
3-methylbutanol
3-methyl-2-buten-1-ol
1-penten-3-ol
1-octen-3-ol
E2-hexenol
Z2-hexenol
E3-hexenol
Z3-hexenol
2,3-butanediol
ethyl acetate
propyl acetate
butyl acetate
pentyl acetate
hexyl acetate
isobutyl acetate
With APL
Without APL
Distance

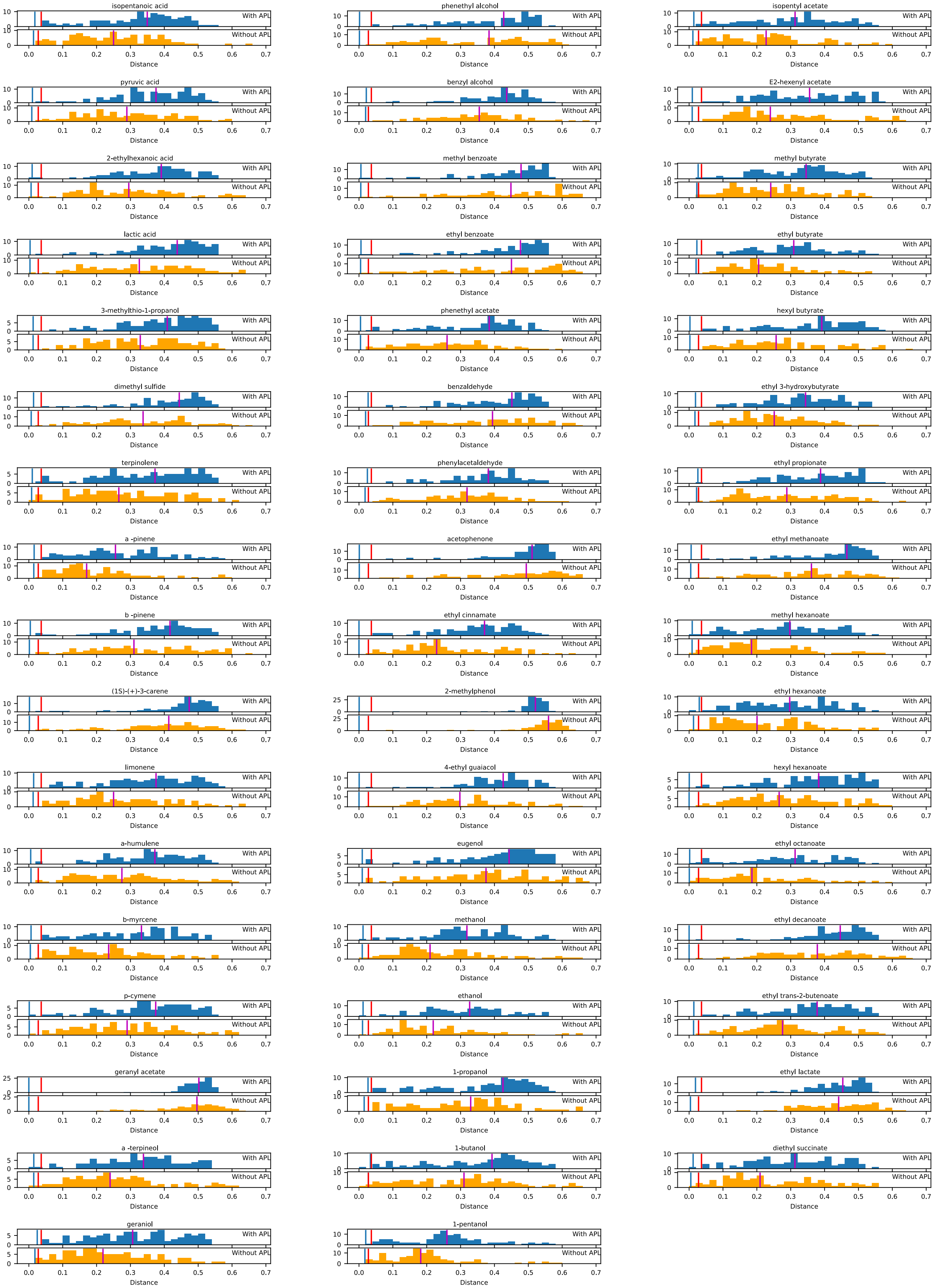


**Figure S2:** Histograms of distances between the marked first spike sequence code of the odorant on top of each panel and the other 103 odorants. Blue vertical line indicates the maximum distance between the marked first spike sequence codes responding to the same odorant at different concentration levels. Top: KC responses of the Calyx circuit model with the APL feedback neuron. Bottom: KC responses of the Calyx circuit model without the APL feedback neuron.
